# Multicolor fluorescence microscopy for surgical guidance using a chip-scale imager with a low-NA fiber optic plate and a multi-bandpass interference filter

**DOI:** 10.1101/2023.10.16.562247

**Authors:** Micah Roschelle, Rozhan Rabbani, Efthymios Papageorgiou, Hui Zhang, Matthew Cooperberg, Bradley A. Stohr, Ali Niknejad, Mekhail Anwar

**Affiliations:** Department of Electrical Engineering and Computer Sciences, University of California, Berkeley, California 94720, USA; Department of Radiation Oncology, University of California, San Francisco, California 94158, USA; Department of Urology, University of California, San Francisco, California 94158, USA; Department of Pathology, University of California, San Francisco, California 94158, USA

## Abstract

In curative-intent cancer surgery, intraoperative fluorescence imaging of both diseased and healthy tissue can help to ensure successful removal of all gross and microscopic disease with minimal damage to neighboring critical structures, such as nerves. Current fluorescence-guided surgery (FGS) systems, however, rely on bulky and rigid optics that incur performance-limiting trade-offs between sensitivity and maneuverability. Moreover, many FGS systems are incapable of multiplexed imaging. As a result, clinical FGS is currently limited to millimeter-scale detection of a single fluorescent target. Here we present a scalable, lens-less fluorescence imaging chip, VISION, capable of sensitive and multiplexed detection within a compact form factor. Central to VISION is a novel optical frontend design combining a low-numerical-aperture fiber optic plate (LNA-FOP) and a multi-bandpass interference filter, which is affixed to a custom CMOS image sensor. The LNA-FOP acts as a planar collimator to improve resolution and compensate for the angle-sensitivity of the interference filter, enabling high-resolution and multiplexed fluorescence imaging without lenses. We show VISION is capable of detecting tumor foci of less than 100 cells at near video framerates and, as proof of principle, can simultaneously visualize both tumor and nerves in *ex vivo* prostate tissue.

## 1. Introduction

A hallmark of a successful curative-intent cancer surgery is the complete removal of all disease, both gross and microscopic, with minimal damage to neighboring healthy tissue. Surgical guidance, however, is largely accomplished through visual and tactile feedback, which lacks adequate contrast and sensitivity, often leading to residual disease in the patient [1]. Thus, positive surgical margins (PSMs)—where cancer cells are detected along the margin of the excised tissue post-operation, suggesting an incomplete resection—occur at significant rates across all common cancers and are particularly prevalent in breast, prostate, and head and neck cancers [2]. In prostate cancer, the focus of this work, PSM rates exceed 20%, the highest PSM rate for men among the ten most common solid organ cancers in the US [2]. Studies show that PSMs are highly correlated to recurrence and mortality and require adjuvant treatment, incurring added costs, toxicity, and burden to the patient [2,3].

Across cancer types, surgeons face a common challenge—without intraoperative imaging to assist with identifying microscopic disease, they are confronted with the difficult decision of empirically removing wide tissue margins that may include undetected disease but risk damage to unseen critical structures. For example, in prostate cancer treatment, lack of visualization results in iatrogenic damage to tumor-adjacent nerves, which remains a persistent problem in radical prostatectomies despite advances in surgical technique [4]. Even in minimally invasive, robot-assisted radical prostatectomies, the prevalence of erectile dysfunction, caused by damage to the cavernous nerves, is posited to range from 10-46% at 1-year post operation [5]. Another challenge lies in the accurate assessment of nodal involvement, which is strongly correlated to aggressive disease and, if detected and removed, can both treat disease and guide adjuvant therapy [6]. Overall, these challenges underscore the need for surgical guidance systems that offer both the molecular-level contrast, sensitivity, and resolution necessary for real-time detection of microscopic disease in addition to multiplexed sensing of diseased and healthy tissue for effective surgical planning.

Among intraoperative imaging modalities, fluorescence imaging is uniquely suited to these needs [7,8]. Targeted fluorescent probes offer molecular specificity and contrast that is straightforward to extend to multiple cellular targets using different color fluorophores. As a result, a variety of contrast agents and fluorescence-guided surgery (FGS) systems are in clinical development [9–11] and fluorescence probes exist for the detection of many common cancer types [9] and, more recently, for critical structures such as nerves [12,13].

While several commercially available FGS systems allow for the clinical application of such agents, these systems suffer from performance-limiting trade-offs between sensitivity and physical maneuverability, as they rely on bulky and rigid optical components. The resolution and collection efficiency (sensitivity) of an optical system are dependent on both the aperture size and proximity to the sample. Highly sensitive microscopic detection, therefore, requires large optics placed near the tissue. Thus, conventional high-resolution systems, such as surgical microscopes [14] (Fig. 1a), employ bulky optics that obstruct the surgical workflow, limit visibility to line-of-sight imaging, and are incompatible with laparoscopic techniques that are standard of care for radical prostatectomies [15]. While fiber optic endoscopes [16,17] offer microscopic detection within a smaller form factor, they are fundamentally limited by tradeoffs between flexibility and field of view (FoV). This can lead to the use of flexible fibers with sub-millimeter FoVs, requiring long scan times to cover the full resection cavity. Finally, laparoscopes (Fig. 1b), seek to enhance maneuverability and visibility by using miniaturized optics that cover a wide FoV [18]. However, miniaturized optics have small apertures that limit sensitivity and resolution. Moreover, to cover and maintain focus across a large and depth-varying surgical FoV, a long working distance and a large depth-of-field—both inversely related to resolution and collection efficiency—are required, resulting in further performance limitations. An extended quantitative discussion of laparoscopes is presented in *supplementary section 1.1*. Thus, while many practical FGS systems are effective at detecting millimeter-scale tumor foci, they are incapable of identifying microscopic disease on the order of hundreds of microns or smaller [19]. Compounded with the absence of multiplexed fluorescence capabilities on many systems [20], this lack of sensitivity prevents conventional FGS systems from taking full advantage of the molecular specificity and multicolor contrast offered by fluorescent probes.

**Fig. 1.**
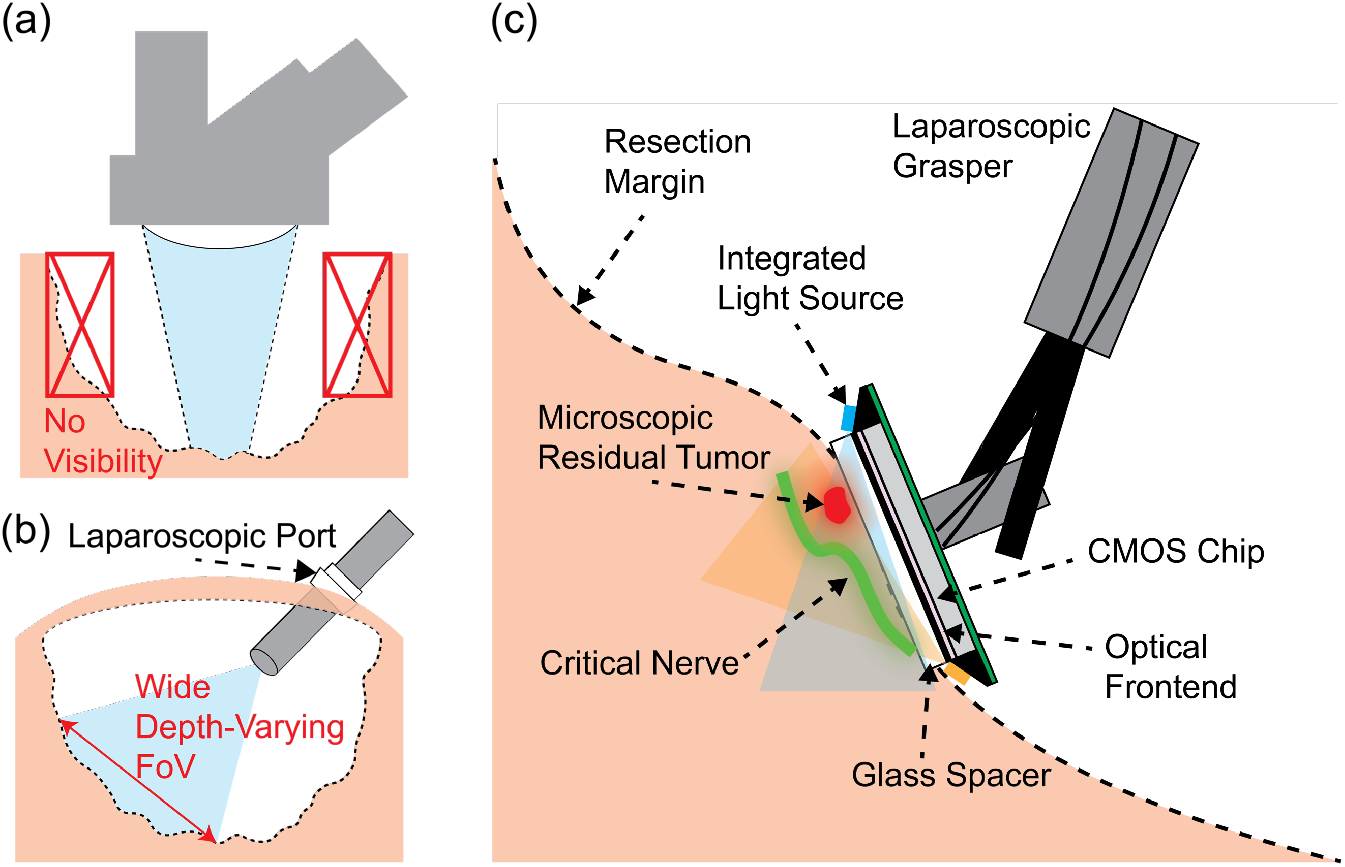
Comparison of conventional FGS systems with VISION. (**a**) Surgical microscopes have limited maneuverability and visibility and are incompatible with minimally invasive surgical techniques. (**b**) Laparoscopes use miniaturized optics that cover a wide depth-varying FoV, resulting in reduced sensitivity. (**c**) Conceptual diagram of VISION. VISION eliminates bulky optical lenses through contact imaging to allow for both a compact form factor and high collection efficiencies necessary for microscopic detection. Moreover, the optical frontend presented in this work enables multicolor fluorescence imaging for simultaneous visualization of both tumor and nerves.

To address these needs in next-generation FGS, we introduce a lens-less and chip-based fluorescence imaging platform, VISION (Versatile Imaging Sensor for Intra-Operative Navigation) (Fig. 1c), capable of sensitive and multiplexed intraoperative imaging within a compact form factor. In contrast to conventional FGS systems, VISION takes advantage of contact imaging to eliminate bulky optical lenses and their associated tradeoffs. In contact imaging, the image sensor is placed in direct contact with the tissue to capture fluorescence emissions before they diverge [21]. This approach allows for adequate resolution without lenses and high collection efficiencies necessary for microscopic detection. Furthermore, CMOS technology enables full sensor integration within a single chip and an inherently array-like structure which can be scaled to accommodate FoVs on the order of 1 cm^2^ without significant trade-offs in resolution [21]. Therefore, the full imaging system can have a planar form factor on the scale of single chip, converting the optical image to an electronic one at the point of imaging, and connected to external hardware through only a few flexible wires. This facilitates eventual integration on the surface of surgical tools and manipulation of the sensor within the resection cavity including to image non-line-of-sight regions.

Despite these advantages, fluorescence contact imaging comes with two unique optical design challenges: (1) integrating high-performance, customizable, multicolor fluorescence emission filters on chip and (2) achieving microscopic resolution without lenses. In this work, we improve on our prior work in chip-based FGS [22–24], introducing a novel optical frontend design that addresses both challenges. Our design consists of a multi-bandpass thin-film interference filter directly coated on a low-numerical-aperture fiber optic plate (LNA-FOP). The LNA-FOP is strongly absorptive of off-axis light and acts as a planar collimator, both to compensate for sensitivity the interference filter and to substantially improve imager resolution by eliminating divergent emissions. This innovation enables the use of high-performance interference filters without conventional focusing optics or lower-performance absorption filters, opening the door to multiplexed fluorescence microscopy on-chip with VISION. Fabricating this optical frontend on our custom-designed CMOS image sensor, we show that VISION achieves a resolution of 110μm and can detect individual cell clusters of less than 100 cells at near video frame rates. Moreover, we image (*ex vivo*) resected patient prostate tissue illustrating three important clinical scenarios in FGS: (1) detection of microscopic disease at the resection margin, (2) simultaneous identification of tumor and nerves, and (3) assessment of spread beyond the primary site.

## 2. Challenges in lens-less fluorescence image sensor design

Chip-based contact imagers require thin, planar alternatives to conventional fluorescence emission filters and focusing optics. Due to the small absorption cross-section and optical properties of conventional fluorophores, fluorescence emissions are often 4 to 6 orders of magnitude weaker than the excitation light in intensity and red-shifted only 10-100nm in wavelength from their absorption peak [25]. As a result, high-performance optical filters are needed for the detection of weak fluorescence signals from an intense excitation background.

### 2.1 Challenges with interference filters for lens-less fluorescence imaging

For conventional microscopes, thin-film interference filters are the gold standard for fluorescence filtering. They can be engineered to have high out-of-band rejection, sharp cut-off transitions, and near-total passband transmittance for any visible or near-IR spectral band as well as multiple non-overlapping passbands for multiplexed imaging [25]. Interference filters, however, are highly sensitive to angle of incidence (AOI) and only maintain performance specifications for normal incident light. At off-axis AOIs, the filter passband shifts to shorter wavelengths, rapidly increasing excitation bleed-through. This property complicates on-chip integration as without collimating lenses it is difficult to ensure that excitation light remains perpendicular to the filter [26]. This is especially true for *in vivo* contact imaging applications where the excitation light must be introduced at an angle between the tissue and the sensor.

### 2.2 Limitations of prior on-chip fluorescence filter designs

As a result, previously published on-chip fluorescence filter designs rely primarily on absorption filters, which are inherently angle-insensitive but have limited design versatility and inferior performance. Absorption filters can be fabricated using a colored dye [27–30] or semiconducting material [31,32] that selectively absorbs excitation light while passing emitted fluorescence. As the absorption spectra is material dependent, different materials and/or fabrication processes must be used to image a new fluorescence wavelength, limiting design flexibility. Moreover, absorption filters have gradual cut-off transitions and significant passband losses that reduce fluorescence collection efficiencies. Finally, dye-based absorption filters often provide inadequate out-of-band rejection due to autofluorescence while semiconductor-based absorption filters are innately long-pass, reducing contrast by passing out-of-band tissue autofluorescence. Hybrid filters combining absorption and interference filters can improve out-of-band rejection, but still retain the poor pass-band characteristics and limited versatility of absorption filters [33,34]. As an alternative, approaches that avoid conventional filters altogether have also been proposed, such as using on-chip nanoplasmonic structures [35] or time-resolved imaging [24,36], but rely on unconventional inorganic fluorophores with either sufficiently long Stokes shifts or long relaxation times, respectively, to achieve adequate rejection. Inorganic fluorophores lack precedence for FDA approval and are often large (10s-1000s of nm), potentially hindering biodistribution [37].

Multicolor fluorescence imaging on chip has proven an even more difficult design challenge. Prior multicolor filter designs [38–41] rely on a hybrid approach stacking both dye-based absorption filters and interference filters to achieve multiple passbands. Such designs require elaborate multi-step fabrication processes and generally suffer from the limitations in versatility and performance inherent to absorption filters.

### 2.3 Resolving optics for lens-less imaging

A second challenge in lens-less imager design is maintaining resolution. Without lenses, resolution is governed by the distance between the sample and the sensor over which incident light can diverge. For practical *in vivo* imaging, this distance cannot be arbitrarily small as a separation distance between the imager and the tissue must be maintained to introduce the excitation light. To improve resolution, planar collimating structures can deblur the image by eliminating divergent emissions at the cost of reduced collection efficiency. These structures can be integrated on-chip as angle-selective gratings (ASGs) fabricated in the metal layers [42,43] or introduced externally with a fiber optic plate [34,39]. Alternatively, coded apertures in combination with computational reconstruction algorithms can be used to improve resolution as well as to recover depth information, but require complex calibration routines, hand-optimized reconstruction algorithms, and constraints on the image sparsity and signal-to-noise ratio (SNR), limiting the robustness of these methods in low-latency *in vivo* applications [38,44,45].

## 3. Materials and Methods

### 3.1 Optical frontend design

Our optical frontend design (Fig. 2a) harnesses the complementary characteristics of a high-performance multi-bandpass interference filter and an LNA-FOP to achieve both versatile multicolor fluorescence filtering and enhanced imager resolution. To compensate for the sensitivity of the interference filter to off-axis excitation, we use the LNA-FOP as a strongly absorptive angle filter. While FOPs and interference filters have been used together in prior on-chip fluorescence filter designs [34,39], these FOPs have NAs close to 1 and are used primarily as a substrate for filter fabrication and to improve imager resolution, not for rejection of off-axis excitation. Instead, in these works, dye-based absorption filters are used in combination with interference filters to provide excitation rejection across all AOIs. Absorption filters introduce less-ideal filter characteristics and limit design versatility such as the incorporation of multiple bandpass regions necessary for multiplexed fluorescence imaging without sacrificing sensitivity through pixel-level patterning of color filters. In this work, the collimating properties of the low-NA FOP eliminate the need for additional absorption filters.

**Fig. 2.**
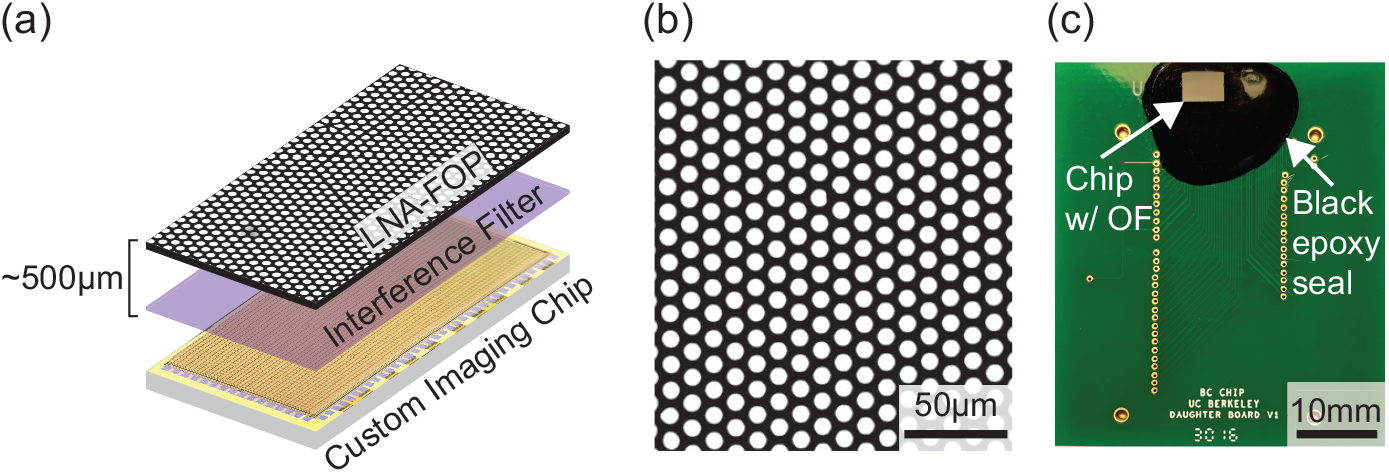
VISION optical frontend design. (**a**) Optical frontend design consisting of a low-NA fiber optic plate (LNA-FOP) and a multi-bandpass interference filter. (**b**) Micrograph of the LNA-FOP. (**c**) Prototype of VISION with the optical frontend (OF) epoxied on top of our custom CMOS image sensor chip mounted on a PCB for testing.

The LNA-FOP (Shenzhen Laser, LTD) has a NA 0.15 and is composed of a matrix of optical fibers embedded in black, absorptive glass (Fig. 2b). It is less than 500μm thick, contains fibers with a 12μm outer diameter and 10μm core, and has 40% transmittance at normal incidence. Light incident within the acceptance angle of the fibers is transmitted by total internal reflection through the fibers. Excitation light incident beyond the acceptance angle, that would otherwise pass through the interference filter, is attenuated in the absorptive sidewalls. As the LNA-FOP absorbs indiscriminate of wavelength, this technique enables the use of any interference filter with single or multiple passbands, allowing for a versatile design adaptable to any visible and NIR fluorophore and, importantly, for multicolor fluorescence imaging. In this work, specifically, we demonstrate dual-color imaging with a dual-bandpass interference filter (ETFITC/Cy5m, Chroma Technology Corp). The LNA-FOP also improves resolution by acting as a collimator.

### 3.2 Custom image sensor and optical frontend fabrication

We fabricate the proposed optical frontend design on our custom CMOS image sensor (Fig. 2c) presented in [22]. The 2.5x5mm chip contains a 4.4x2.2mm image sensor with an 80x36 active pixel array of 44x44μm^2^ pixels at a 55μm pitch. The sensor achieves 8.2 V s^-1^ pW^-1^ sensitivity at 700nm and uses correlated double sampling to attain shot-noise-limited performance in exposure times as short as 50ms.

The optical frontend can be fabricated in two ways: (1) from commercially available parts (for rapid prototyping) including a 250μm LNA-FOP and the dual-bandpass interference filter deposited on a 1mm-thick glass substrate and (2) by directly coating the interference filter on a 500μm-thick LNA-FOP to reduce the overall thickness. In (2) a 500μm-thick FOP is used to minimize the risk of fracturing during deposition, but a 250μm-thick FOP could likely be used with reduced yield. Direct deposition (2) reduces overall thickness and has the additional benefit of reducing imaging artifacts near the edge of the sensor (*supplementary section 1.2*).

The optical component(s) are epoxied directly onto the chip. First, all surfaces are cleaned with 99% isopropyl alcohol. Next, the optical epoxy (SYLGARD 184, Dow Chemicals) is mixed and degassed for 20 minutes in a vacuum chamber. The optical component(s) are then epoxied on the chip and cured in a controlled temperature chamber (45 minutes at 65°C). To prevent optical leakage and protect the bond wires, the edges of the imager are sealed with black epoxy (EP1046FG Black, Resinlab), cured at 85°C for 90 minutes.

Due to scattering through the FOP, the order that the FOP and interference filter are placed on the chip effects filter performance under different illumination scenarios. Following the reasoning in *supplementary section 1.3*, in this work, the filter is placed below the FOP to block scattered excitation under oblique illumination.

### 3.3 Preparation of cell cultures

PC3-PIP cells, which overexpress prostate-specific membrane antigen (PSMA), are cultured on monolayer plates. To form sparse clusters of closely spaced cells, the cells are trypsinized from monolayer cultures and plated on a commercially available matrix. Cells from the matrix cultures were fixed onto glass slides with care to ensure that they remain in small (10-1000 cell) clusters. Separate sets of slides are stained for imaging in each fluorescent channel with an anti-PSMA rabbit primary antibody (D4S1F, Cell Signaling Technology) and either an IRDye680LT goat anti-rabbit (LI-COR) or an AlexaFluor488 goat anti-rabbit secondary antibody (Invitrogen).

### 3.4 Preparation of prostate tissue samples

Banked resected patient prostate tissue samples are obtained from UCSF (IRB 15-16090). The paraffin-embedded tissue blocks are sectioned at 4μm and mounted onto glass slides. One representative slide from each block is stained with hemoxylin and eosin (H&E) for histological analysis. The remaining slides are used for immunofluorescence staining. To label prostate cancer cells, an anti-PSMA rabbit primary antibody (D4S1F, Cell Signaling Technology) is used with an IRDye680LT goat anti-rabbit secondary antibody (LI-COR). For nerve labeling, an anti-S100 mouse primary (4C4.9, Abcam) is used with an AlexaFluor488 anti-mouse secondary (Invitrogen).

### 3.5 Imaging with VISION

A custom printed circuit board (PCB) is used to supply power and bias currents for the imaging chip as well as to read out and digitize each captured image frame. All timing and control signals necessary for image acquisition are generated by a field-programmable gate array (XEM6010, Opal Kelly), which interfaces between the chip and a laptop. A custom software user interface is used to set the imaging parameters and to visualize and save image data.

Two fiber-coupled lasers with wavelengths at 488nm and 633nm (QFLD-488-20S and QFLD-633-20S, QPhotonics) are used for fluorescence excitation. They are driven by a TEC-controlled driver (6310 Combo Source, Arroyo Instruments) and are operated with an optical intensity of 28mW/cm^2^ for imaging. The lasers are collimated and transilluminate the sample at approximately 70°. While transillumination is difficult *in vivo*, a glass separator or light guide plate can be used in future work to couple light introduced from the side for epi-illumination between the sensor and tissue [22,46]. In this work, a 1mm-thick glass separator is fixed on top of the sensor to accurately reflect the reduction in resolution and sensitivity with the added thickness of such a device.

All imaging is performed in a light-proof box to minimize background light. For routine imaging, the sample is placed directly on the surface of the sensor and excitation light is introduced through the back of the slide. To simultaneously image the same FoV with VISION and a benchtop fluorescence microscope, we use an inverted microscope (Leica DM-IRB). The sample slide is placed with the coverslip facing away from the microscope objective lens. VISION is inverted and placed on the opposite side of the slide as the microscope objective. Excitation light is introduced at an angle between the objective and slide.

For each region of interest (ROI), 100 images are acquired with VISION at 50-75ms exposure time to be used for image analysis or averaging. Signal-to-noise ratio (SNR) is reported as the maximum single pixel SNR within the ROI and is calculated as the mean pixel value divided by the standard deviation of the pixel intensity across 100 images. Image post-processing including thresholding and false coloring is performed in ImageJ (NIH).

## 4. Results and Discussion

### 4.1 Optical frontend characterization

Fig. 3a shows the measured angular transmittance of the 250μm-thick LNA-FOP in air at 488nm compared with that of the on-chip angle-selective gratings (ASGs) presented in our prior work [22]. Angular transmittance measurements were performed using an optical power meter (PM100D with S120C photodiode, ThorLabs) and the same fiber-coupled lasers used for imaging. Narrow-band interference filters for each laser line (ZET488/10x and ET645/30x, Chroma Technologies Corp) eliminate any out-of-band emissions from the lasers. The sample is clamped into a motor-controlled rotating stage (HDR50, ThorLabs) to measure the transmitted optical power at 1° increments.

**Fig. 3.**
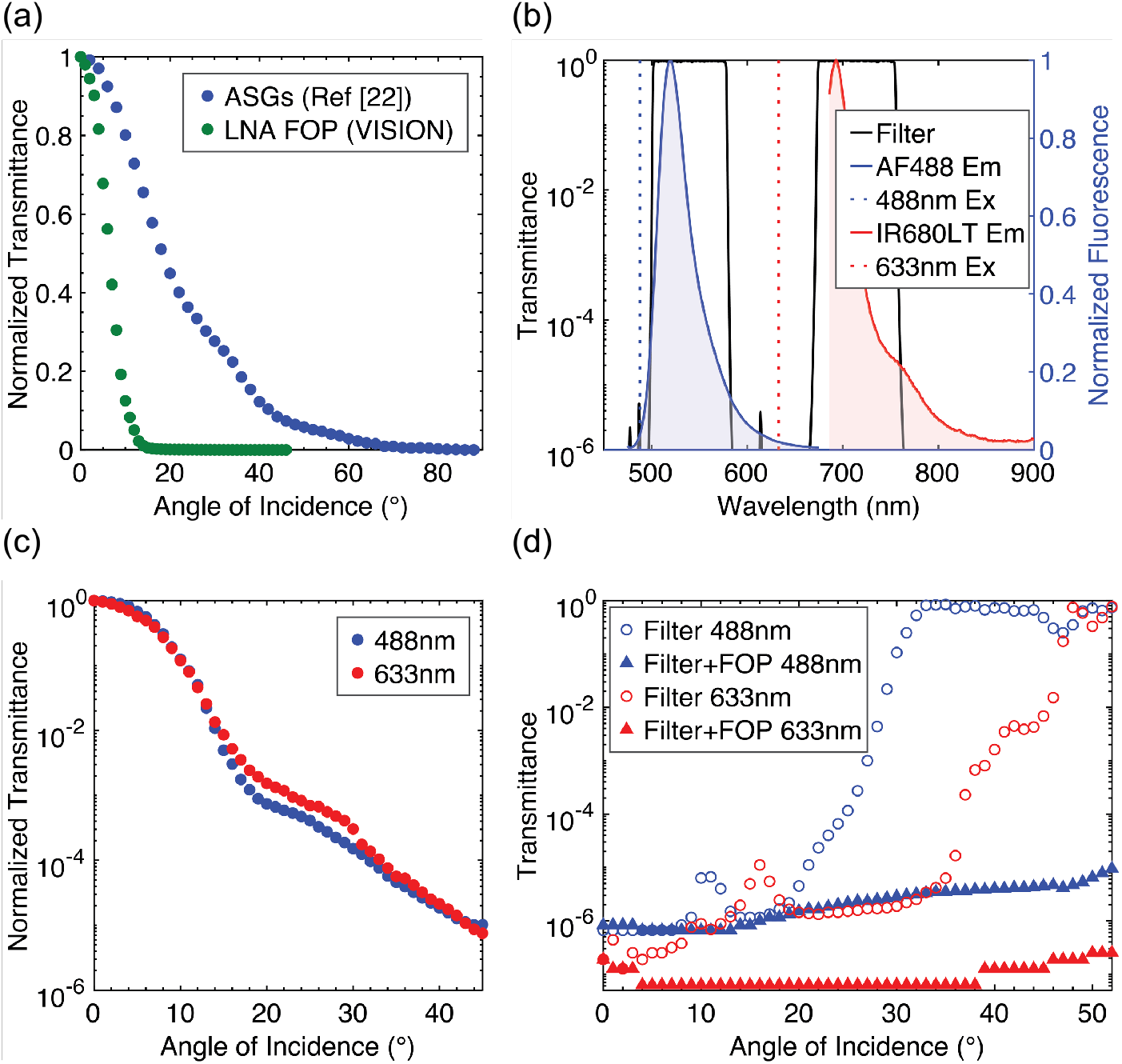
Optical frontend characterization. (**a**) Measured angular transmittance of the LNA-FOP compared with that of the on-chip angle-selective gratings (ASGs) presented in our prior work [22]. (**b**) Transmittance spectra of the dual-bandpass interference filter. (**c**) Angular transmittance of the LNA-FOP at 488nm and 633nm plotted on a log scale. (**d**) Angular transmittance of the 488nm and 633nm excitation lasers through both the interference filter alone and the interference filter with the LNA-FOP.

Compared with the ASGs, that are fabricated using reflective metal and have aspect ratios restricted by CMOS process design rules, the LNA-FOP achieves significantly sharper angle-selectivity and stronger rejection of oblique light with minimal added thickness. Accordingly, the measured full width at half maximum (FWHM) of the FOP is 12.65° in air, almost 3x narrower than that of the ASGs (36°). For a lens-less imager with collimators, the FWHM of the collimator is directly related to the resolution of the system.

Fig. 3b shows the normal incidence transmittance spectra of the dual-bandpass interference filter (data is reproduced with permission from Chroma Technologies Corp). The filter has less than 10^−6^ transmittance at both 488nm and 633nm, the excitation wavelengths for AlexaFluor488 (AF488) and IRDye680LT (IR680LT) which are the fluorophores selected for our imaging studies. Moreover, the sharp band-edge and high passband transmittance (>95%) of the filter allow for efficient collection of fluorescence emissions from each fluorophore. However, as noted previously, excitation bleed-through increases sharply with AOI. Angular transmittance measurements of the filter at both excitation wavelengths (Fig. 3d) show a roughly 100x increase in transmittance at 24° for 488nm and 36° for 633nm. By 33° and 48°, respectively, the filter is practically transparent to the excitation, passing nearly all incident light.

We compensate for this behavior with the LNA-FOP, which is strongly absorptive of off-axis light. Fig. 3c shows the measured angular transmittance of the FOP on a log scale. At 45°, the FOP achieves less than 10^−5^ transmittance of both wavelengths. The rejection increases for AOIs larger than 45°. Therefore, as shown in Fig. 3d, when both the FOP and interference filter are used together, the resulting optical frontend maintains less than 10^−5^ transmittance at 488nm and less than 10^−6^ transmittance at 633nm across all AOIs.

To illustrate the potential performance improvements over absorption filters gained through the improved passband characteristics of the interference filter, we compare the VISION optical frontend with the amorphous silicon (a-Si) absorption filter presented in our prior work [32]. As discussed in *supplementary section 1.4*, we measure a 9.3x improvement in excitation rejection and a 4.3x improvement in fluorescence emission collection efficiency with the new design.

### 4.2 Resolution measurements

To measure the resolution of VISION, we image a standard USAF-1951 resolution test target as shown in Fig. 4a. For fluorescence imaging, a negative USAF target patterns excitation light onto a layer of Cy5.5 dye (Sulfo-Cyanine5.5 NHS ester, Abcam) dissolved in DMSO at 100μg/mL. The dye is contained on the target with a 150μm-thick glass coverslip and is placed on VISION for imaging. A 1mm-thick glass separator separates the sensor from the sample. For practical *in vivo* imaging, the glass separator would be used to introduce excitation light to the tissue.

**Fig. 4.**
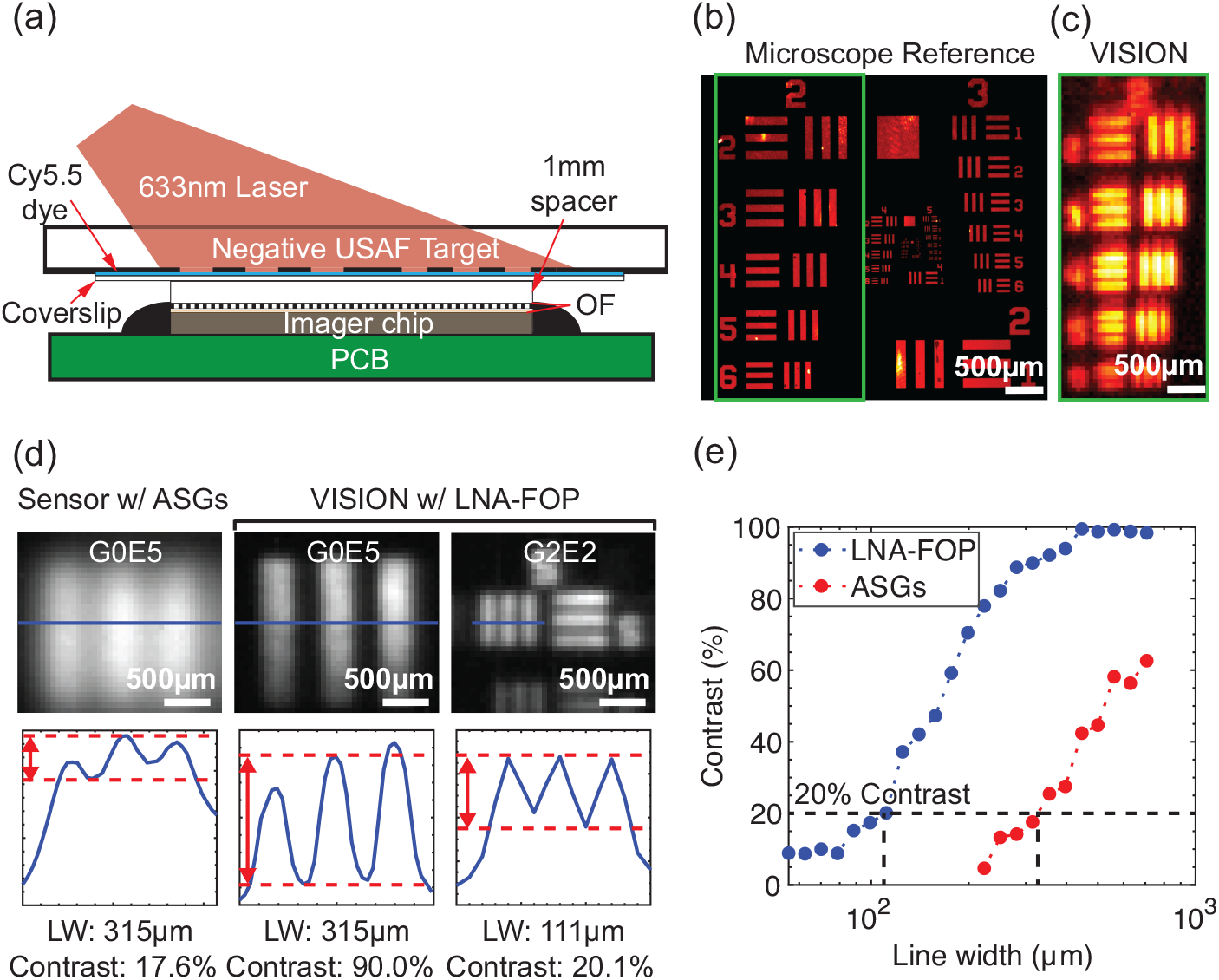
Resolution measurements. (**a**) Experimental set up for resolution measurements. (**b**) Reference image of the fluorescent test target from a benchtop fluorescence microscope using a 2.5x objective. Green box is around group 2. (**c**) Image taken with VISION of group 2 on the test target, which spans VISION’s resolution limit. (**d**) Comparison of representative images of individual 3-bar elements from the target taken with a sensor with the ASGs presented in our prior work [22] and with VISION. (**e**) Plot of the measured contrast of each 3-bar element as function of line width from images taken with VISION and the sensor with ASGs.

A reference image of the target taken with a benchtop fluorescence microscope (Fig. 4b) can be compared qualitatively to an image taken with VISION of group 2 on the target (Fig. 4c), which spans the resolution limit of our system. By group 2, element 6 (line widths of 70 μm), the lines become nearly indistinguishable with VISION.

To quantify the resolution improvement achieved with the LNA-FOP, we image individual 3-bar elements on the target both with VISION and an imager with ASGs presented in our prior work [22] and measure the image contrast. As the line widths approach the resolution limit of the sensor, the blur reduces the contrast between light and dark bars. Contrast is calculated as

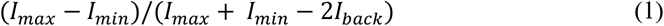

where *I*_*max*_ is the maximum pixel value in the bright bars on the target, *I*_*min*_ is the minimum pixel value in the dark bars, and *I*_*back*_ is the background (the average pixel value when the excitation source is off). Fig. 4d shows representative images from this measurement along with the line scans through each element used to calculate contrast.

The plot in Fig. 4e shows measured image contrast as a function of line width for both imagers. These measurements show that with the LNA-FOP, VISION can resolve line widths of 110μm at 20% contrast, while with the ASGs, the imager is only capable of line widths of 330μm at 20% contrast, representing a roughly 3-fold improvement in resolution. As expected, this result corresponds with the approximately 3x difference in FWHM between the LNA-FOP and the ASGs shown in Fig. 3a. The resolution can be further improved by reducing the 1mm separation distance between the sample and imager, but some separation must be preserved to introduce the excitation light to the tissue.

To be sure, this improvement in resolution gained with the increased angle selectivity of the FOP comes with a reduction in fluorescence signal seen by the sensor. However, with increased resolution, the background at each pixel due to tissue autofluorescence or nonspecific staining is also reduced improving the achievable signal-to-background ratio (SBR) for tumor foci smaller than the resolution of the system [19,47]. In FGS, the SBR often plays a dominant role in detection.

### 4.3 Sensitivity measurements

The identification and removal of microscopic tumor foci (<1mm^3^ or <10^5^-10^6^ cells) remains a critical barrier to the success of curative-intent cancer surgeries. Imaging of microscopic foci requires both efficient collection of fluorescence emissions and sufficient resolution to achieve high SNRs and signal to background ratios necessary for detection [19,47]. Towards these ends, VISION utilizes a contact imaging approach to achieve high collection efficiencies for microscopic detection.

We quantify the sensitivity of VISION by imaging slides containing differently sized clusters of fluorescently labeled PC3-PIP prostate cancer cells. The slides are imaged using excitation intensities of 28mW/cm^2^ and exposure times of 50ms and 75ms for the AF488 and IR680LT channels, respectively. Fig. 5a.i shows a representative image of a AF488-stained sample taken with VISION compared with a reference image from benchtop fluorescence microscope (Fig. 5a.ii). From 20x microscope images (Fig. 5b.i-ii), the cell clusters labeled A and B are determined to contain approximately 100 and 50 cells, respectively. These clusters are imaged on VISION with maximum single pixel SNRs of 22x (13dB) and 10x (10dB), respectively.

**Fig. 5.**
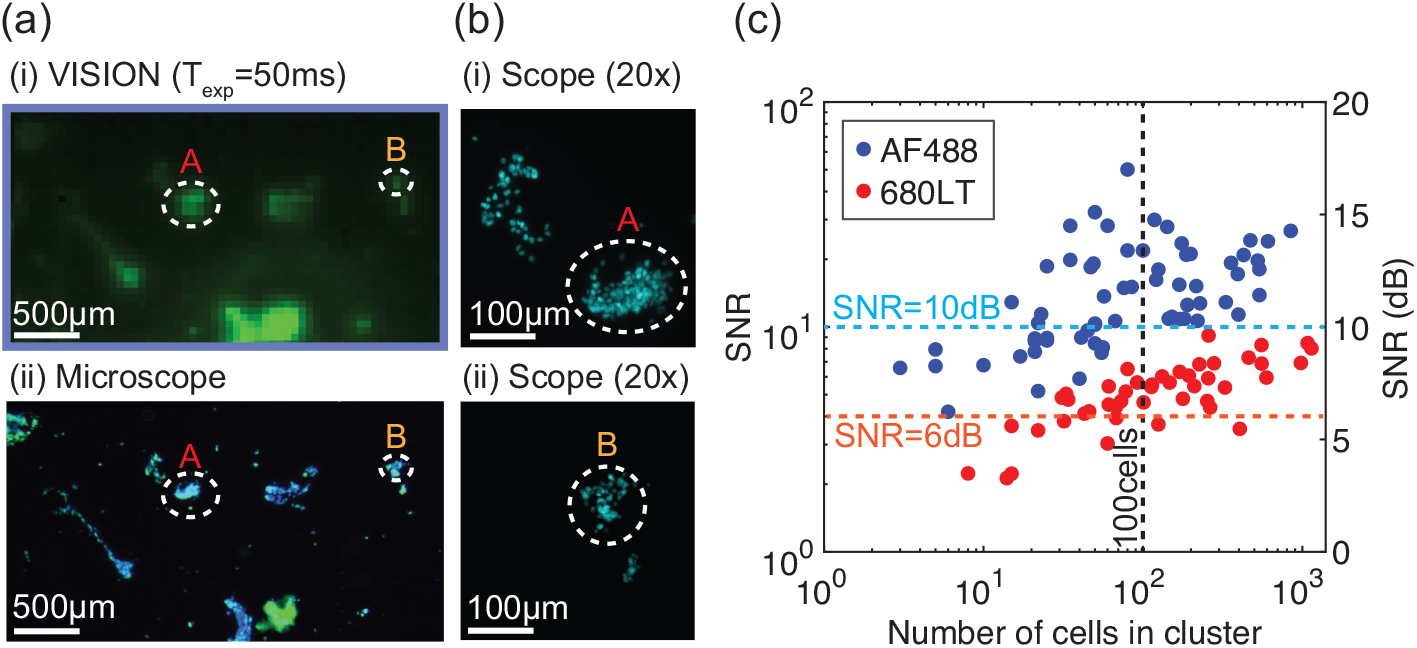
Sensitivity measurements. (**a**) (i) A representative image taken with VISION and (ii) a microscope reference of the PC3-PIP cell culture slides fluorescently stained with AF488. Clusters marked with A and B contain ∼100 and ∼50 cells and are imaged on VISION with SNRs of 22x and 10x, respectively. (**b**) 20x microscope images of clusters (i) A and (ii) B used for counting the number of cells in each cluster. (**c**) Plot of measured maximum single pixel SNR vs. cluster size for PC3-PIP cell cultures stained with AF488 and IR680LT.

This procedure is repeated for a set of samples containing clusters ranging in size from 1-1000 cells. Fig. 5c shows a plot of the maximum measured single-pixel SNR versus the cell cluster size for each fluorescence channel. VISION can detect cell clusters as small as 100 cells in a single 50ms or 75ms exposure with SNRs greater than 10x (10dB) and 4x (6dB) in the AF488 and IR680LT channels, respectively. Cells stained with AF488 are inherently brighter due to higher quantum yield of the dye and excitation closer to the absorption peak. Sensitivity can be further increased through longer exposure times, higher excitation intensities, and hardware optimizations (see Conclusion) advancing VISION closer to single-cell detection.

### 4.4 Ex vivo prostate tissue imaging

To demonstrate imaging in clinically relevant scenarios in prostate cancer resection with VISION, we image *ex vivo* banked, resected patient prostate tissue interspersed with tumor and nerves (Fig. 6). The tissue is fluorescently stained for both prostate cancer with an anti-PSMA-IR680LT (red) and nerves anti-S100-AF488 (green). While anti-S100 is an intracellular probe and, therefore, is not suitable for *in vivo* imaging, it is used here as a proof-of-principle for multiplexed imaging and can be replaced by existing nerve-specific intraoperative probes [12,13] in future experiments. To verify staining, slides are scanned with a fluorescence microscope (Zeiss Axio Observer) and compared against full-slide H&E scans by a trained pathologist (Fig. 6a.iii-iv).

**Fig. 6.**
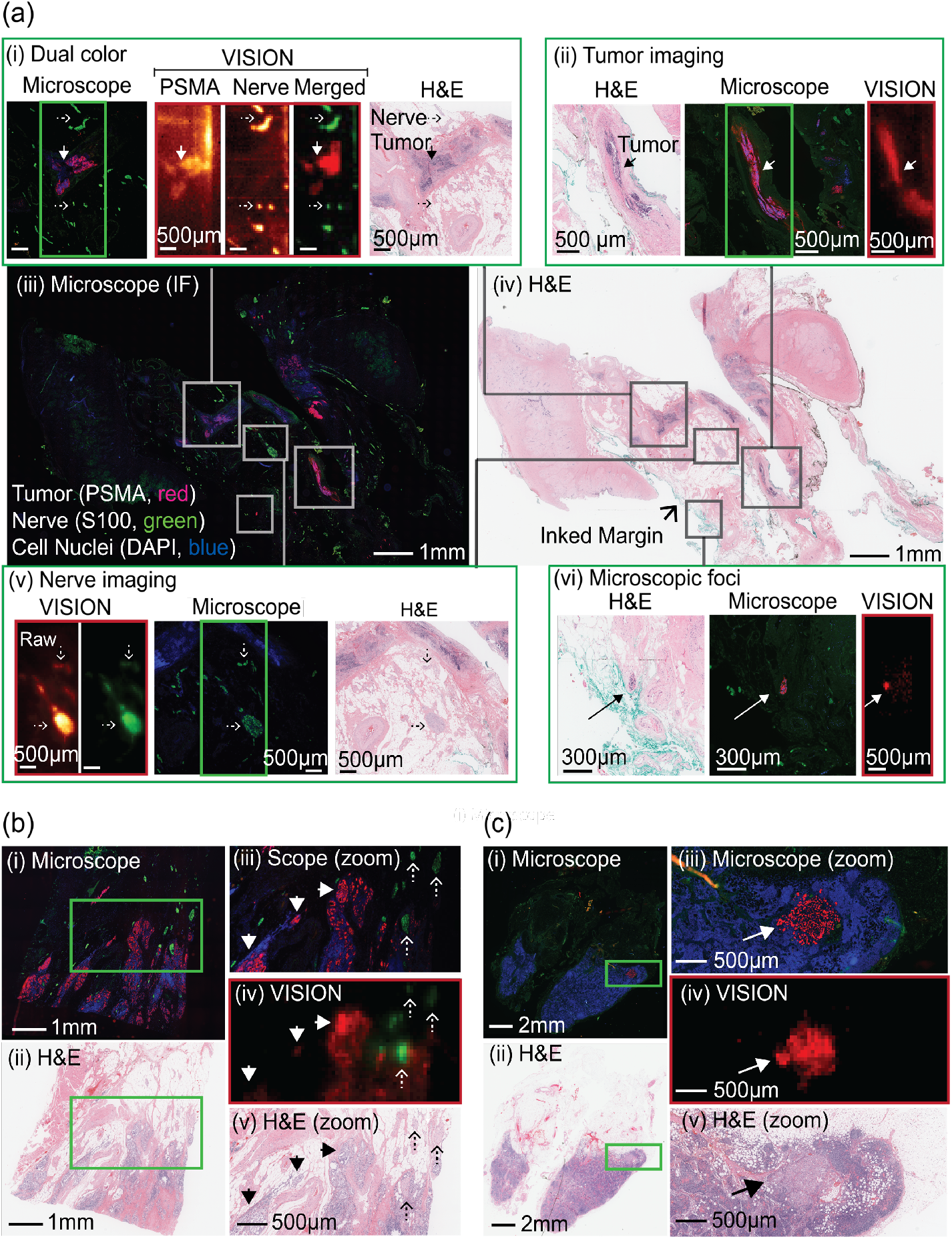
*Ex vivo* imaging of resected prostate tissue with tumor foci and nerves. (**a**) (iii & iv) Full-slide microscope scans of immunofluorescence (IF) and H&E stained slides of resected prostate tissue. (i) Simultaneous detection of both tumors and nerves with VISION is possible by overlaying separate exposures of each channel. (ii & v) VISION can clearly identify major tumor and nerves in the sample with microscopic resolution. (vi) VISION detects a microscopic focus (<100 cells) along the inked margin. (**b**) Images of extra-prostatic extension into fibroadipose tissue. (i) IF and (ii) H&E full slide scans of sample with ROI for imaging in green. (iv) VISION detects tumor foci and nerves present in (iii) IF and (v) H&E references. (**c**) Images of metastatic tumor in a lymph node. (i) IF and (ii) H&E full slide scans of sample with ROI for imaging in green. (iv) VISION detects a small focus of metastatic tumor visible in (iii) IF and (v) H&E references.

Relevant areas of each slide are then imaged in each fluorescence channel with VISION. Exposure times of 50ms and 75ms are used for the AF488 and IRDye680LT channels, respectively. To best demonstrate the resolution and image accuracy of VISION, the SNR of the images is enhanced by 10dB by averaging 100 individual frames. While this level of averaging is used for presentation of the results and, in practice, would result in long imaging times, a single frame is necessary for detection of the labeled cells within the samples. To this point, single frame maximum at-pixel SNRs are reported in the following section for each image and are within the range of those measured with the PC3-PIP cell cultures. In addition, Fig. S4 compares single frame images with the averaged images presented in Figs. 6a.ii and 6a.v and illustrates the increase in SNR with averaging.

Fig. 6a.i-ii and 6a.v-vi show a side-by-side comparison of images captured with VISION and a benchtop fluorescence microscope alongside H&E images for an histological reference. As shown in Fig 6a.i, dual-color imaging of both channels is achieved with our dual-bandpass filter by taking a separate exposure with each laser and then overlaying the images. In Figs. 6a.ii and 6a.v, VISION identifies major clusters of both tumor cells (SNR=9.3dB) and nerves (SNR=10-25.5dB), respectively. To emphasize the sensitivity of VISION, Fig. 6a.vi shows VISION detecting a microscopic tumor focus of less than 100 cells (SNR=7.3dB) located along the inked margin of the sample. Figs. 6b and 6c illustrate scenarios in which VISION can detect and characterize tumor spread beyond the primary tumor site. In Fig 6b, VISION visualizes extra-prostatic extension of tumor into fibroadipose tissue interspersed with nerves and, in Fig. 6c, VISION detects a small lymph node metastasis (SNR=8.8dB). Overall, these images illustrate the potential use of VISION to intraoperatively assess resection margins for microscopic residual disease, provide critical information regarding the spatial proximity of tumor and nerves for surgical guidance, and identify potential sites of metastatic spread.

It should be noted that line-like artifacts are visible along the edges of some images (especially in Fig. 6a.i). As discussed in *supplementary section 1.2*, these artifacts occur when using the optical frontend design with the commercial interference filter and are caused by excitation light reflected off the bond-pads of the sensor and guided through the 1mm-thick fused silica substrate. The artifacts can be eliminated by directly depositing the filter on the FOP. As a result of these artifacts as well as the lower collection efficiency and use of trans-illumination compared with the microscope, the background signal due to the excitation bleed-through is more pronounced in the VISION images.

To be sure, after successful demonstration of *ex vivo* imaging, there are still challenges to address before transitioning to *in vivo* imaging. The cell cultures used for sensitivity measurements in this study represent an idealized scenario in which there is no background signal from surrounding tissue. Additionally, *in vivo* targets may be obscured by thin layer of blood or tissue, which attenuate signals through scattering and absorption. For these reasons, poorer sensitivity and reduced SNRs may be encountered *in vivo*, but can be compensated for through the strategies discussed in the preceding section. On the other hand, the cell cultures contain no more than 4 layers of cells and the tissue sections are less than a single layer thick (∼4μm). In contrast, *in vivo* sub-millimeter tumor foci are 3D structures consisting of tens to hundreds of layers of cells, possibly leading to more concentrated signals.

## 5. Conclusion

Fluorescence-guided surgery technologies show potential to significantly improve clinical outcomes in curative-intent cancer surgery by helping surgeons better detect and eliminate residual and microscopic disease while avoiding damage to neighboring critical tissues. However, current FGS systems using conventional optics have restricted maneuverability, are limited to macro-scale detection, and for the most part are not designed for multiplexed imaging. Therefore, we have developed VISION, a fluorescence contact-imaging chip capable of both multiplexed and microscopic detection without sacrificing maneuverability. Specifically, VISION takes advantage of lens-less contact imaging and CMOS technology to: (1) achieve efficient collection of fluorescence emissions and resolution necessary for microscopic detection; (2) allow for a scalable design for imaging FoVs up to centimeter dimensions with little trade-off in resolution; and (3) enable a compact and planar form factor for eventual integration with laparoscopic tools and that can be manipulated to image surfaces throughout the resection cavity.

Moreover, in this work, we demonstrated a lens-less optical frontend combining an interference filter and LNA-FOP to provide both high-performance multicolor fluorescence filtering and enhanced resolution. We showed that the LNA-FOP effectively compensates for the angle-sensitivity of the filter, achieving less than 10^−5^ transmittance of excitation light across all AOIs. Critical to lens-less imaging applications, our design is thin, planar, and inherently scalable in size. By coating the filter directly on the FOP, the total thickness is just over 500μm. Most importantly, compared with prior work—which primarily focused on absorption filters— our optical frontend harnesses the inherent advantages of interference filters: sharp cut-offs and near 100% passband transmittance, a versatile design adaptable for any visible or near-infrared fluorophore, and easy extension to multicolor imaging with organic fluorophores. While we demonstrated dual-color fluorescence imaging, three- and four-band interference filters are widely available such that our design can be straightforwardly adapted for complex multicolor fluorescence imaging schemes. The LNA-FOP also improves imager resolution. With the 250μm LNA-FOP, VISION can resolve linewidths of 110μm with 20% contrast at 1mm separation distance from the sample, representing a 3x improvement over our prior work.

The advantages of our optical frontend design have relevance beyond intraoperative imaging. Other applications of fluorescence contact imaging include miniaturized implantable imagers for functional brain imaging in free-moving animals [36,48,49] and for *in vivo* monitoring of therapeutic response [50,51] as well as compact lab-on-chip devices for high-throughput molecular screening and diagnostics [52–55]. In particular, the versatility of our optical frontend allows for transferrable sensor design: adapting a sensor to a new fluorescence wavelength involves epoxying on a new interference filter and changing the excitation source. We believe this design flexibility can help to bring about new applications which leverage novel optical probes and multiplexed imaging techniques.

Relevant to intraoperative imaging, we showed that VISION can detect microscopic tumor foci of less than 100 cells with SNRs near or above 10dB at exposure times of 50ms, close to acceptable video frame rates. These results show the potential for VISION to help surgeons identify microscopic residual disease below the mm-scale sensitivity limit of currently available intraoperative imaging platforms. Moreover, to emphasize clinical relevance for FGS, we demonstrated that VISION, enabled by the optical frontend presented in this work, can simultaneously identify microscopic tumor foci and nerves in *ex vivo* resected prostate tissue. We illustrated three important clinical scenarios in which VISION can improve surgical outcomes: (1) detection of a microscopic tumor foci of less than 100 cells along the surgical margin, (2) simultaneous localization of residual tumor and nerves, and (3) identification of a metastatic lymph node.

Future improvements to VISION can further this potential. Sensitivity can be increased through hardware optimizations. First, the optical frontend can be optimized with a custom interference filter with optimal passbands for the fluorophores. The FWHM of the FOP can also be increased to improve collection efficiency (albeit, at the cost of resolution) while still compensating for the angle-sensitivity of the interference filter which can maintain rejection up to 20-35°. Second, the quantum efficiency of our photodiodes (currently 13% at 530nm and 18% at 700nm) can be enhanced through use of a more optimized optical CMOS process. Building on our proof-of-concept platform, VISION can be adapted for practical use in the clinic. While our prototype sensor has a FoV 4.4x2mm, CMOS image sensors are inherently scalable [56], enabling larger FoVs with the same resolution (pixel size) for faster raster scanning over the surgical cavity. In the future, for example, the sensor size could be increased to 1x1cm, imaging a 1cm^2^ area in <1s while still fitting through a 12mm laparoscopic port. In addition, CMOS technology enables the integration of power management, control, readout, and image sensor circuits all on a single mm-scale chip. Thus, a future system including the PCB and housing can be integrated at the scale of the chip and connected to external hardware only by a few flexible wires as shown in the conceptual Fig. 1c. Finally, a new multi-bandpass filter can be integrated in the optical frontend to extend VISION to tri-color imaging with multiple NIR bands at 700nm and 800nm to cover a wide array of available FGS probes [9].

In conclusion, this work shows and furthers the promise of chip-based fluorescence contact imagers, such as VISION, in filling in the performance gaps left by conventional lens-based FGS systems in terms of maneuverability, sensitivity, and multiplexed imaging. With these advantages and further improvements, we envision chip-based intraoperative imagers like VISION improving clinical outcomes by extending surgical vision to hard-to-access areas and to microscopic and multiplexed imaging of both diseased and healthy tissue.

## Acknowledgements

Additional support for this work was provided by philanthropic contributions from the John V. Carbone Jr. Pancreas Cancer Research Memorial Fund. The authors thank Chroma Technologies Corp. for assistance with custom filter depositions and the sponsors of BSAC (Berkeley Sensors and Actuators Center) and TSMC for chip fabrication.

## Data availability

Data underlying the results presented in this paper are not publicly available at this time but may be obtained from the authors upon reasonable request.

## Supplemental Document

See Supplement 1 for supporting content.

## References

1. K. R. Tringale, J. Pang, and Q. T. Nguyen, “Image-guided surgery in cancer: A strategy to reduce incidence of positive surgical margins,” Wiley Interdiscip Rev Syst Biol Med 10, (2018).

2. R. K. Orosco, V. J. Tapia, J. A. Califano, B. Clary, E. E. W. Cohen, C. Kane, S. M. Lippman, K. Messer, A. Molinolo, J. D. Murphy, J. Pang, A. Sacco, K. R. Tringale, A. Wallace, and Q. T. Nguyen, “Positive Surgical Margins in the 10 Most Common Solid Cancers,” Sci Rep 8, (2018).

3. J. L. Wright, B. L. Dalkin, L. D. True, W. J. Ellis, J. L. Stanford, P. H. Lange, and D. W. Lin, “Positive Surgical Margins at Radical Prostatectomy Predict Prostate Cancer Specific Mortality,” Journal of Urology 183, 2213–2218 (2010).

4. A. Kumar, V. R. Patel, S. Panaiyadiyan, K. R. Seetharam Bhat, M. C. Moschovas, and B. Nayak, “Nerve-sparing robot-assisted radical prostatectomy: Current perspectives,” Asian J Urol 8, 2–13 (2021).

5. T. F. N. Lima, J. Bitran, F. S. Frech, and R. Ramasamy, “Prevalence of post-prostatectomy erectile dysfunction and a review of the recommended therapeutic modalities,” Int J Impot Res 33, 401–409 (2021).

6. L. Cheng, H. Zincke, M. L. Blute, E. J. Bergstralh, B. Scherer, and D. G. Bostwick, “Risk of prostate carcinoma death in patients with lymph node metastasis,” Cancer 91, 66–73 (2001).

7. I. S. Alam, I. Steinberg, O. Vermesh, N. S. van den Berg, E. L. Rosenthal, G. M. van Dam, V. Ntziachristos, S. S. Gambhir, S. Hernot, and S. Rogalla, “Emerging Intraoperative Imaging Modalities to Improve Surgical Precision,” Mol Imaging Biol 20, 705–715 (2018).

8. F. J. Voskuil, J. Vonk, B. van der Vegt, S. Kruijff, V. Ntziachristos, P. J. van der Zaag, M. J. H. Witjes, and G. M. van Dam, “Intraoperative imaging in pathology-assisted surgery,” Nat Biomed Eng (2021).

9. C. W. Barth and S. Gibbs, “Fluorescence image-guided surgery: a perspective on contrast agent development,” in (SPIE-Intl Soc Optical Eng, 2020), p. 18.

10. B. Zhu and E. M. Sevick-Muraca, “A review of performance of near-infrared fluorescence imaging devices used in clinical studies,” British Journal of Radiology 88, (2015).

11. T. Nagaya, Y. A. Nakamura, P. L. Choyke, and H. Kobayashi, “Fluorescence-guided surgery,” Front Oncol 7, (2017).

12. L. G. Wang, C. W. Barth, C. H. Kitts, M. D. Mebrat, A. R. Montaño, B. J. House, M. E. Mccoy, A. L. Antaris, S. N. Galvis, I. Mcdowall, J. M. Sorger, and S. L. Gibbs, I M A G I N G Near-Infrared Nerve-Binding Fluorophores for Buried Nerve Tissue Imaging (2020), Vol. 12.

13. L. G. Wang, C. W. Barth, C. H. Kitts, M. D. Mebrat, A. R. Montaño, B. J. House, M. E. McCoy, A. L. Antaris, S. N. Galvis, I. McDowall, J. M. Sorger, and S. L. Gibbs, “Near-infrared nerve-binding fluorophores for buried nerve tissue imaging,” Sci Transl Med 12, eaay0712 (2020).

14. L. Ma and B. Fei, “Comprehensive review of surgical microscopes: technology development and medical applications,” J Biomed Opt 26, (2021).

15. A. J. Costello, “Considering the role of radical prostatectomy in 21st century prostate cancer care,” Nat Rev Urol 17, 177–188 (2020).

16. G. Walker, A. P. Wang, P. Z. McVeigh, Z. Ivanishvilli, W. Siu, E. J. Seibel, H. Lesiuk, F. Alkherayf, and B. J. Drake, “The Scanning Fiber Endoscope: A Novel Surgical and High-Resolution Imaging Device for Intracranial Neurosurgery,” Operative Neurosurgery 23, 326–333 (2022).

17. D. Shin, M. C. Pierce, A. M. Gillenwater, M. D. Williams, and R. R. Richards-Kortum, “A fiber-optic fluorescence microscope using a consumer-grade digital camera for in vivo cellular imaging,” PLoS One 5, (2010).

18. A. V Dsouza, H. Lin, E. R. Henderson, K. S. Samkoe, and B. W. Pogue, “Review of fluorescence guided surgery systems: identification of key performance capabilities beyond indocyanine green imaging,” J. Biomed. Opt 21, 80901 (2016).

19. B. W. Pogue, “Perspective review of what is needed for molecular-specific fluorescence-guided surgery,” J Biomed Opt 23, 1 (2018).

20. F. van Beurden, D. M. van Willigen, B. Vojnovic, M. N. van Oosterom, O. R. Brouwer, H. G. van der Poel, H. Kobayashi, F. W. B. van Leeuwen, and T. Buckle, “Multi-Wavelength Fluorescence in Image-Guided Surgery, Clinical Feasibility and Future Perspectives,” Mol Imaging 19, (2020).

21. A. Greenbaum, W. Luo, T. W. Su, Z. Göröcs, L. Xue, S. O. Isikman, A. F. Coskun, O. Mudanyali, and A. Ozcan, “Imaging without lenses: Achievements and remaining challenges of wide-field on-chip microscopy,” Nat Methods 9, 889–895 (2012).

22. E. P. Papageorgiou, B. E. Boser, and M. Anwar, “Chip-Scale Angle-Selective Imager for in Vivo Microscopic Cancer Detection,” IEEE Trans Biomed Circuits Syst 14, 91–103 (2020).

23. H. Najafiaghdam, C. C. S. Pedroso, N. A. Torquato, B. E. Cohen, and M. Anwar, “Fully Integrated Ultra-thin Intraoperative Micro-imager for Cancer Detection Using Upconverting Nanoparticles,” Mol Imaging Biol (2022).

24. H. Najafiaghdam, E. Papageorgiou, N. A. Torquato, B. Tian, B. E. Cohen, and M. Anwar, “A 25 micron-thin microscope for imaging upconverting nanoparticles with NIR-I and NIR-II illumination,” Theranostics 9, 8239–8252 (2019).

25. J. Reichman, Handbook of Optical Filters for Fluorescence Microscopy (2000).

26. M. Dandin, P. Abshire, and E. Smela, “Optical filtering technologies for integrated fluorescence sensors,” Lab Chip 7, 955–977 (2007).

27. E. Carregal-Romero, C. Fernández-Sánchez, A. Eguizabal, S. Demming, S. Büttgenbach, A. Llobera, M. L. Chabinyc, D. T. Chiu, J. C. McDonald, A. D. Stroock, J. F. Christian, A. M. Karger, A. Cornwell, S. Beecher, A. Raja, D. D. Bradley, A. J. Demello, J. C. Demello, A. Llobera, S. Demming, H. N. Joensson, J. Vila-Planas, H. Andersson-Svahn, and S. Büttgenbach, Development and Integration of Xerogel Polymeric Absorbance Micro-Filters into Lab-on-Chip Systems (2004), Vol. 96.

28. E. Yıldırım, Ç. Arpali, and S. A. Arpali, “Implementation and characterization of an absorption filter for on-chip fluorescent imaging,” Sens Actuators B Chem 242, 318–323 (2017).

29. O. Hofmann, X. Wang, A. Cornwell, S. Beecher, A. Raja, D. D. C. Bradley, A. J. DeMello, and J. C. DeMello, “Monolithically integrated dye-doped PDMS long-pass filters for disposable on-chip fluorescence detection,” Lab Chip 6, 981–987 (2006).

30. Y. Dattner and O. Yadid-Pecht, “Low light cmos contact imager with an integrated poly-acrylic emission filter for fluorescence detection,” Sensors 10, 5014–5027 (2010).

31. J. Alex Chediak, Z. Luo, J. Seo, N. Cheung, L. P. Lee, and T. D. Sands, “Heterogeneous integration of CdS filters with GaN LEDs for fluorescence detection microsystems,” in Sensors and Actuators, A: Physical (2004), Vol. 111, pp. 1–7.

32. E. P. Papageorgiou, H. Zhang, B. E. Boser, C. Park, and M. Anwar, “Angle-insensitive amorphous silicon optical filter for fluorescence contact imaging,” Opt Lett 43, 354 (2018).

33. C. Richard, A. Renaudin, V. Aimez, and P. G. Charette, “An integrated hybrid interference and absorption filter for fluorescence detection in lab-on-a-chip devices,” Lab Chip 9, 1371–1376 (2009).

34. K. Sasagawa, A. Kimura, M. Haruta, T. Noda, T. Tokuda, and J. Ohta, “Highly sensitive lens-free fluorescence imaging device enabled by a complementary combination of interference and absorption filters,” Biomed Opt Express 9, 4329 (2018).

35. L. Hong, H. Li, H. Yang, and K. Sengupta, “Fully Integrated Fluorescence Biosensors On-Chip Employing Multi-Functional Nanoplasmonic Optical Structures in CMOS,” IEEE J Solid-State Circuits 52, 2388–2406 (2017).

36. J. Choi, A. J. Taal, W. L. Meng, E. H. Pollmann, J. W. Stanton, C. Lee, S. Moazeni, L. C. Moreaux, M. L. Roukes, and K. L. Shepard, “Fully Integrated Time-Gated 3D Fluorescence Imager for Deep Neural Imaging,” IEEE Trans Biomed Circuits Syst 14, 636–645 (2020).

37. S. Bouccara, G. Sitbon, A. Fragola, V. Loriette, N. Lequeux, and T. Pons, “Enhancing fluorescence in vivo imaging using inorganic nanoprobes,” (n.d.).

38. A. J. Taal, C. Lee, J. Choi, B. Hellenkamp, and K. L. Shepard, “Toward implantable devices for angle-sensitive, lens-less, multifluorescent, single-photon lifetime imaging in the brain using Fabry–Perot and absorptive color filters,” Light Sci Appl 11, (2022).

39. N. Kulmala, K. Sasagawa, T. Treepetchkul, H. Takehara, M. Haruta, H. Tashiro, and J. Ohta, “Lensless dual-color fluorescence imaging device using hybrid filter,” Jpn J Appl Phys 61, (2022).

40. W. S. Hee, K. Sasagawa, A. Kameyama, A. Kimura, M. Haruta, T. Tokuda, and J. Ohta, “Lens-free dual-color fluorescent CMOS image sensor for förster resonance energy transfer imaging,” Sensors and Materials 31, 2579–2594 (2019).

41. T. Treepetchkul, N. Kulmala, K. Sasagawa, H. Takehara, M. Haruta, H. Tashiro, and J. Ohta, Dual-Color Lensless Fluorescence Imaging by Using a Notch Interference Filter and Absorption Filters (2021).

42. E. P. Papageorgiou, B. E. Boser, and M. Anwar, “An angle-selective CMOS imager with on-chip micro-collimators for blur reduction in near-field cell imaging,” in Proceedings of the IEEE International Conference on Micro Electro Mechanical Systems (MEMS) (Institute of Electrical and Electronics Engineers Inc., 2016), Vol. 2016-February, pp. 337–340.

43. K. Sugie, K. Sasagawa, M. C. Guinto, M. Haruta, T. Tokuda, and J. Ohta, “Implantable CMOS image sensor with incident-angle-selective pixels,” Electron Lett 55, 729–731 (2019).

44. J. K. Adams, D. Yan, J. Wu, V. Boominathan, S. Gao, A. V. Rodriguez, S. Kim, J. Carns, R. Richards-Kortum, C. Kemere, A. Veeraraghavan, and J. T. Robinson, “In vivo lensless microscopy via a phase mask generating diffraction patterns with high-contrast contours,” Nat Biomed Eng (2022).

45. S. Moazeni, E. H. Pollmann, V. Boominathan, F. A. Cardoso, J. T. Robinson, A. Veeraraghavan, and K. L. Shepard, “A Mechanically Flexible Implantable Neural Interface for Computational Imaging and Optogenetic Stimulation over 5.4 × 5.4 mm 2FoV,” in Digest of Technical Papers - IEEE International Solid-State Circuits Conference (Institute of Electrical and Electronics Engineers Inc., 2021), Vol. 64, pp. 288–290.

46. H. Shin, G.-W. Yoon, W. Choi, D. Lee, H. Choi, D. S. Jo, N. Choi, J.-B. Yoon, and I.-J. Cho, “Miniaturized multicolor fluorescence imaging system integrated with a PDMS light-guide plate for biomedical investigation,” npj Flexible Electronics 7, 7 (2023).

47. A. Gharia, E. P. Papageorgiou, S. Giverts, C. Park, and M. Anwar, “Signal to Noise Ratio as a Cross-Platform Metric for Intraoperative Fluorescence Imaging,” Mol Imaging 19, (2020).

48. R. Rebusi, J. P. Olorocisimo, J. Briones, Y. Ohta, M. Haruta, H. Takehara, H. Tashiro, K. Sasagawa, and J. Ohta, “Simultaneous CMOS-Based Imaging of Calcium Signaling of the Central Amygdala and the Dorsal Raphe Nucleus During Nociception in Freely Moving Mice,” Front Neurosci 15, (2021).

49. Y. Sunaga, Y. Ohta, Y. M. Akay, J. Ohta, and M. Akay, “Monitoring neural activities in the VTA in response to nicotine intake using a novel implantable microimaging device,” IEEE Access 8, 68013–68020 (2020).

50. R. Rabbani, H. Najafiaghdam, B. Zhao, M. Zeng, V. Stojanovic, R. Muller, and M. Anwar, “A 36×40 Wireless Fluorescence Image Sensor for Real-Time Microscopy in Cancer Therapy,” in Proceedings of the Custom Integrated Circuits Conference (Institute of Electrical and Electronics Engineers Inc., 2022), Vol. 2022-April.

51. R. Rabbani, H. Najafiaghdam, M. M. Ghanbari, E. P. Papageorgiou, B. Zhao, M. Roschelle, V. Stojanovic, R. Muller, and M. Anwar, “Towards an Implantable Fluorescence Image Sensor for Real-Time Monitoring of Immune Response in Cancer Therapy,” in 2021 43rd Annual International Conference of the IEEE Engineering in Medicine & Biology Society (EMBC) (IEEE, 2021), pp. 7399–7403.

52. R. R. Singh, D. Ho, A. Nilchi, G. Gulak, P. Yau, and R. Genov, “A CMOS/Thin-Film Fluorescence Contact Imaging Microsystem for DNA Analysis,” IEEE Transactions on Circuits and Systems I: Regular Papers 57, 1029–1038 (2010).

53. J. Balsam, M. Ossandon, Y. Kostov, H. A. Bruck, and A. Rasooly, “Lensless CCD-based fluorometer using a micromachined optical Söller collimator,” Lab Chip 11, 941–949 (2011).

54. H. Takehara, M. Nagasaki, K. Sasagawa, H. Takehara, T. Noda, T. Tokuda, and J. Ohta, “Micro-light-pipe array with an excitation attenuation filter for lensless digital enzyme-linked immunosorbent assay,” in Japanese Journal of Applied Physics (Japan Society of Applied Physics, 2016), Vol. 55.

55. J. Ohta, H. Takehara, K. Sasagawa, H. Takehara, T. Noda, and T. Tokuda, “On-chip fluorescence detection system with high-density microchamber array based on CMOS image sensor,” in Proceedings - IEEE International Symposium on Circuits and Systems (Institute of Electrical and Electronics Engineers Inc., 2016), Vol. 2016-July, pp. 2867–2870.

56. R. Turchetta, N. Guerrini, and I. Sedgwick, “Large area CMOS image sensors,” Journal of Instrumentation 6, C01099–C01099 (2011).

