## Supplemental document 1 for "Multicolor fluorescence microscopy for surgical guidance using a chip-scale imager with a low-NA fiber optic plate and a multi-bandpass interference filter"

#### *1.1 Quantitative analysis of laparoscopic fluorescence imagers compared with VISION*

Fluorescence laparoscopes such as Intuitive Surgical's da Vinci Xi Firefly utilize miniaturized optics covering a wide FoV to image the full resection cavity with a rigid optical system that fits through 8 or 12mm laparoscopic port. These design choices significantly reduce the achievable sensitivity with these systems. This section provides a light throughput calculation considering measured dimensions from the da Vinci Xi Firefly to compare the sensitivity of these types of systems with VISION.

The key parameter in determining resolution and sensitivity of a lensed imaging system is the numerical aperture (NA), which is a measure of the range of angles over which an optical system can collect light. The NA, is defined as

$$NA = n \sin(\theta) \approx \frac{n\,r}{WD} \tag{1}$$

where $n$ is the index of refraction of the media surrounding the imager and $\theta$ is half angle of the maximum cone of light that can enter or exit the lens. Under the paraxial approximation ($\theta$ is small), the relation simplifies, where $r$ is the radius the of aperture and $WD$ is the working distance, the distance along the optical axis from the lens to the object plane. We measure that the 12mm da Vinci Xi Firefly has an approximate 2mm lens aperture for each of its binocular cameras. According to a surgeon (Matthew Cooperberg, co-author) the device is operated a closest working distance of 2cm and is typically operated at a distance of 5-10cm in order to cover a wide enough FoV. Thus, the NA of the system varies from 0.02-0.1.

The sensitivity of an imaging system is determined by the amount of available light collected from the sample and is quantified by the collection efficiency. For a low-NA system, the collection efficiency can be found geometrically from the solid angle subtended by the lens,

$$CE \approx \frac{NA^2}{4} \tag{2}$$

Thus, based purely on our mechanical measurements, we expect the CE of the Firefly to be between 0.01-0.25%.

The diffraction-limited resolution of an optical system is also dependent on the numerical aperture. For a diffraction-limited system of a given NA imaging a self-luminous sample, the Rayleigh resolution is,

$$\delta = \frac{0.61\,\lambda}{NA} \tag{3}$$

where $\lambda$ is the wavelength of the emitted light. Assuming a wavelength of 700nm (near the peak emission of IRdye680LT used in this work), the expected resolution of the Firefly is between 4.27-21.3µm.

However, the system covers a wide topologically varying FoV and must maintain focus across this FoV, requiring a large depth of field (DoF), which drives the need for a lower resolution. For a low-NA system, the DoF can be calculated as

$$DoF = \frac{n\,\lambda}{NA^2} + \frac{n}{NA} \cdot \frac{e}{M} \approx \frac{n\,\lambda}{NA^2} \tag{4}$$

Where the first term is due to diffraction limited resolution and the second is due to the effective size of the pixel taking into account magnification, $\frac{e}{M}$. For a diffraction-limited system, the above approximation can be made. Thus, if the Firefly were diffraction limited, the DoF would range between 70μm-1.75mm at 2cm and 10cm working distances, respectively. However, displacement of the prostate due to respiration and normal patient movement during surgery has been report to exceed 1cm [1] and tissue variation across a FoV of multiple centimeters could have similar variation. Therefore, it is reasonable to expect a 1cm depth of focus is required. In fact, other reported wide-field fluorescence imaging systems such as the Fluobeam and Solaris have reported DoFs on the order of 1cm with lateral resolutions of approximately 100μm [2]. According to equations (4) and (2), to get a resolution of 100μm with a DoF of 1cm requires a numerical aperture of 0.01 resulting in a collection efficiency of 0.0025%. This result illustrates how larger DoF requirements drive lower lateral resolution and NAs, reducing collection efficiency of lens-based systems.

For comparison, in this work, the collection efficiency of VISION taking into account the angular transmittance and insertion loss of the FOP, is 0.18%. In our prior work [3] with the ASGs, which are less restrictive of angle, the collection efficiency is calculated to be 2.13%. As noted in the Conclusion of this work, the FOP can be optimized to have a higher NA while still compensating for the angle sensitivity of the interference filter in order to improve the collection efficiency of VISION. Based on these calculations, it is reasonable to expect around an order of magnitude improvement in collection efficiency or sensitivity with an optimized version of VISION compared to laparoscopic imager. The difference could be larger depending on the particular specifications of the laparoscope such as the DoF. It is likely that a lens-based system would have resolution at least as good as the 110μm achieved with VISION. However, microscopic detection is primarily driven by SNR and does not require microscopic resolution [4]. To be sure, DoF is not of the same concern with the contact imagers as the imager is placed in direct contact with the tissue minimizing the effect of motion and the FoV is typically smaller than $1cm^2$.

#### *1.2 Mitigating imaging artifacts with the VISION optical frontend*

Line artifacts, such as those shown in Fig. S2b, appear on VISION when the optical frontend is constructed with a commercial interference filter due to excitation light that is reflected off of the on-chip bond pads and wires and passes through the glass filter substrate. As in Fig. S2a, bond pads are located along the bottom and right edges of our custom CMOS image sensor. Bond wires provide electrical connection between the on-chip pads and pads on the carrier PCB to supply power to the chip as well as to interface important control and data lines with the FPGA. To prevent damage to these delicate bond wires, the optical frontend must be epoxied only over the region of the chip occupied by the image sensor, leaving the pads and wire bonds exposed.

Fig. S2b shows an image of the bleed-through from the 633nm excitation laser taken with VISION when using the optical frontend incorporating a commercial filter. The image is taken with an exposure of 50ms and the same laser power used for imaging and is averaged 100 times to accentuate the artifacts. It should be noted that the artifacts appear along the edges of the sensor where the bond pads are located. While the artifacts are relatively low in intensity (approximately 4.5x as bright as the mean sensor noise for a single 50ms exposure), these artifacts affect the image quality when imaging dim or microscopic samples.

Fig. S2d illustrates how the artifacts are produced when VISION is packaged with an optical frontend that uses a commercial interference filter. Most commercial interference filters are deposited on glass substrates that are typically 1mm thick. Since the interference filter must be placed below the FOP to block the scattered component of the excitation (see Fig. S3), this filter substrate separates the FOP from the interference filter. Note that the interference filter is placed in contact with the chip to minimize optical leakage through the sides of the structure. When taking an image, some excitation light scatters through the FOP and reaches the wire bonds and bond pads, which are made of gold and are highly reflective. Some of these rays are reflected in such a way that they are guided by reflection through the filter substrate. If the excitation light is incident at an angle such that it can pass through the interference filter it will

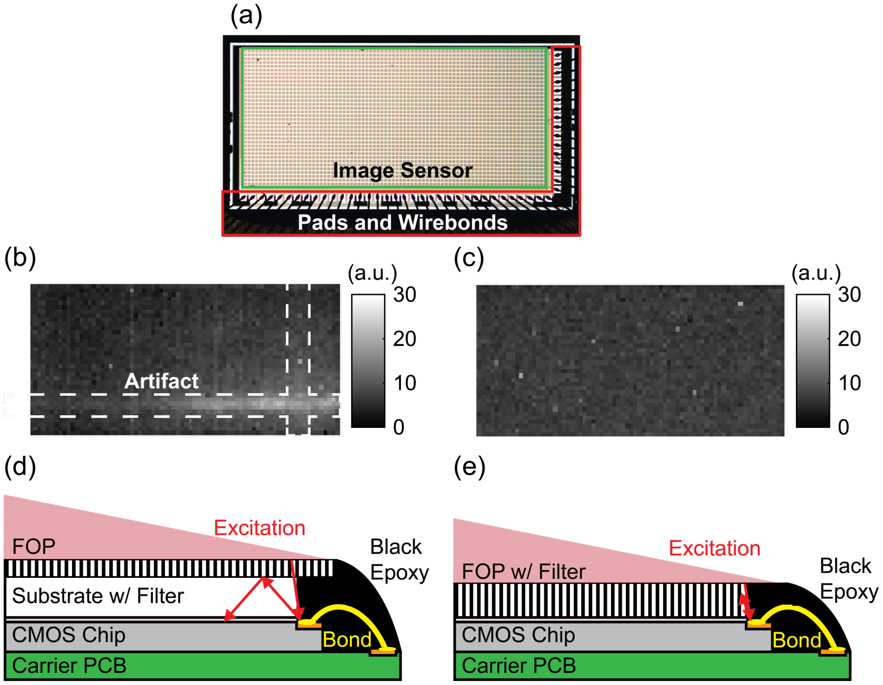


**Fig. S2. Mitigating imaging artifacts.** (**a**) The image sensor chip is electrically connected to the carrier PCB through wire bonds located along the bottom and right sides of the sensor. (**b**) Image of 633nm excitation light taken with imager using the commercial interference filter showing line-like artifacts along the edges of the sensor where the bond pads are present. (**c**) A similar image taken with a imager using the optical frontend with the interference filter directly deposited on the FOP. Note that the artifacts are no longer present. (**d**) These artifacts occur when the commercial interference filter is used due to the 1mm-thick glass substrate that creates a gap between the FOP and the sensor through which excitation light reflected off the bond pads can travel. (**e**) The glass substrate and, thus, the artifacts can be eliminated by depositing the interference filter directly on the FOP.

reach the sensor and produce an artifact. The artifact appears away from the edges of the image because the filter is still able to reject scattered excitation with AOIs close to normal incidence. As a result, the artifacts appear further away from the edges of the image for 633nm excitation than for 488nm excitation as the filter is capable of rejecting excitation across a broader set of AOIs because 633nm is further from the band-edge of the filter than 488nm (see Fig. 3d in the main text).

As illustrated in Fig. S2e, the artifacts can be eliminated by using a custom optical frontend in which the FOP is used directly as the substrate for filter deposition. When fabricated in this way, there is no need for the additional glass substrate separating the FOP from the interference filter. Therefore, excitation light that is reflected off of the bond pads must pass through the absorptive sidewalls of the FOP and is attenuated before reaching the sensor. To be sure, this design includes a more complex fabrication process, but is relatively simple given that the FOP is composed of glass and is compatible with many deposition techniques.

Fig. S2c shows an image of the 633nm laser excitation bleed through taken with VISION using an optical frontend in which the interference filter is directly deposited on a 0.5mm-thick FOP. The line artifact is no longer present and only a slight excitation background registers in the image.

#### *1.3 Orientation dependence of the optical frontend*

The order in which the fiber optic plate (FOP) and interference filter are stacked on the imager has a significant effect on filter performance depending on whether oblique or near-normal incident excitation is used. This dependence is due to the combination of the angular sensitivity of the interference filter and scattering from the FOP. Scattering in the FOP is caused both by material imperfections and by diffraction effects due to the micron-scale fiber apertures.

First, consider the case when excitation is near normal incidence. In this case, the excitation light incident on the FOP passes through the fibers. As the light exits the fibers, a small fraction of the light is scattered to larger angles. Consequently, if the FOP is placed on top of the filter (Fig. S3a.i), excitation light scattered at large AOIs will transmit directly through the filter as described in the preceding section. This effect can be minimized by placing the interference filter on top of the FOP (Fig. S3a.ii). Since the interference filter provides strong excitation rejection near normal incidence, an insignificant amount of excitation light will reach the sensor. Therefore, *for excitation near normal incidence the interference filter is primarily responsible for providing excitation rejection and should always be placed on top of the FOP*.

Now consider the case when the excitation light is incident with a large AOI. When excitation light is incident on the FOP with AOIs larger than the acceptance angle of the fibers, the light will pass through the absorptive sidewalls and experience significant attenuation. However, a small fraction of the incident light scatters through the apertures, escaping the absorptive side walls, and exiting the FOP at near-normal incidence to the sensor. Consequently it is necessary to place the interference filter below the FOP to block the excitation that is scattered. If the interference filter is placed above the FOP, the excitation rejection will be limited by the scattering effects and not the absorption performance of the FOP. Thus, *for obliquely incident excitation the FOP is primarily responsible for providing excitation rejection and should always be placed on top of the interference filter*.

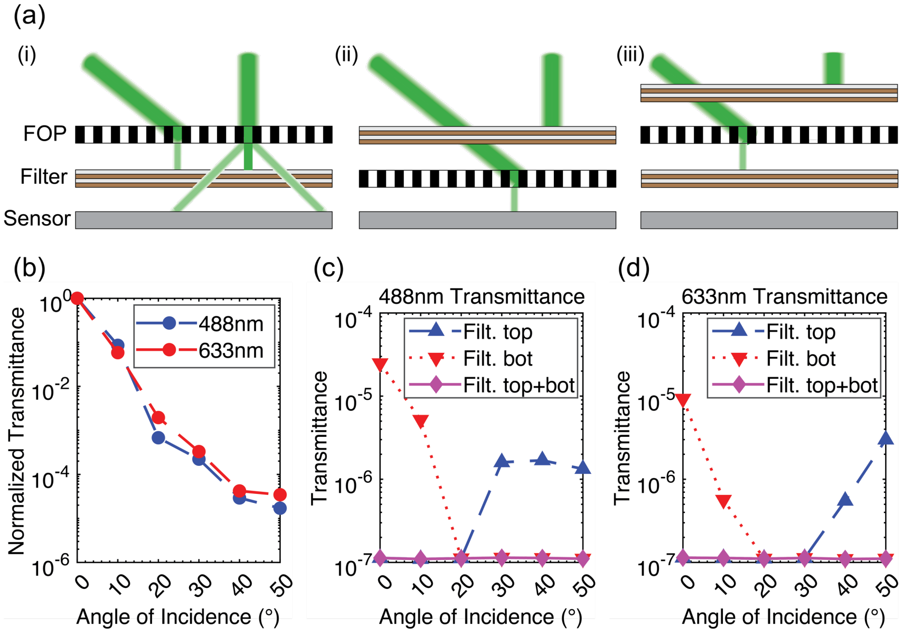


**Fig. S3. Orientation dependence of the VISION optical frontend.** (**a**) (i) When the FOP is placed above the filter, there is strong rejection of obliquely incident excitation, but for normal incident excitation, light scattered through the FOP passes through the interference filter and reaches the sensor. (ii) On the other hand, when the filter is placed above the FOP, normal incident excitation is adequately rejected, but oblique excitation passes through the filter and small amount is scattered through the FOP to the sensor. (iii) For the maximum excitation rejection across all AOIs, the same interference filter can be placed on both sides of the FOP to minimize scattering effects in both cases. (**b**) Measurements of the angular transmittance of the FOP at both excitation wavelengths, show that the transmittance plateaus near 40° due to scattering effects. (**c, d**) Angular transmittance measurements of the optical frontend illustrate the orientation dependent effects described in (a) for both excitation wavelengths.

Measurements of the angular transmittance of the FOP at 488nm and 633nm (Fig. S3b), show that the transmittance of the FOP starts to plateau around 40° at $10^{-5}$ due to scattering effects. To include scattered light in the measurement, the FOP is placed in direct contact with the photometer such that all transmitted light is captured. Similar measurements are performed for both orientations of the optical frontend and the case in which the interference filter is placed on both sides (Fig. S3cd). As explained in the preceding section, the optical frontend best blocks excitation near normal incidence when the interference filter is on top. Conversely, it shows the highest performance for oblique excitation with the FOP on top. The difference in performance between the two orientations at the extremes of both regimes is between 1-2 orders of magnitude, illustrating why the correct choice of orientation is critical. The measurements also show that the angle-sensitivity of the frontend can be mitigated by placing the same interference on both sides of the FOP. With this modification the filter shows superior performance for both normal incident and oblique excitation. Of course this design comes at the cost of added fabrication complexity and is only necessary when the AOI of the excitation light is not known. As discussed in the main text, for contact imaging applications, the excitation must be introduced obliquely, so it is sufficient to have a single interference filter on the bottom of the FOP.

#### *1.4 Comparison of the VISION optical frontend with a-Si absorption filters*

To demonstrate the performance improvements over absorption filters gained through this approach, we compare the proposed optical frontend with the 15μm-thick amorphous silicon (a-Si) filter presented in our prior work [5].

As observed in the measured optical transmittance spectra of both filters (Fig. S4a), the interference filter exhibits a more ideal filter characteristic with a sharper band-edge and higher pass-band transmittance, which maximizes collection of fluorescence emissions. The total collected fluorescence emission is dependent on the collection efficiency of the optics, the absorption of the dye at the chosen excitation wavelength, and the overlap of the dye emission spectra with the filter passband. For the a-Si filter, the gradual roll-off in the filter response forces a significant trade-off between efficient excitation of the fluorophore and fluorescence emission collection. When the a-Si filter is used with IRDye680LT, which has a Stoke's shift of 19nm, similar to many conventional fluorophores, the laser excitation must be at 633nm for adequate excitation rejection, which is only at 33% of the absorption peak. At the emission peak (693nm), the passband transmittance of the 15μm-thick a-Si filter is just 1.6% and rises to maximum pass-band transmittance of 54% due to reflection losses from the high refractive index (4.3) of the a-Si. On the other hand, with the proposed optical frontend, the laser excitation wavelength can be as close to 10nm from the filter band-edge without significant excitation bleed-through (as demonstrated in main Fig. 3d with the 488nm laser excitation), allowing for both optimal excitation of the fluorophore and fluorescence emission collection. This fact is clear in the effective emission spectra of IRDye680LT for both filters (Fig. S4b), which is product of the dye emission spectra and the filter transmittance. The total fraction of emitted fluorescence passed by the a-Si filter is 4.24x less than that passed by the interference filter. Further improvements in the fluorescence collection efficiency of the proposed optical frontend can be made by choosing an interference filter with a band-edge closer to the absorption edge of the dye to allow for excitation of 680LT at the absorption peak (676nm).

These results are confirmed by sensitivity measurements performed by imaging a 2x serial dilution of IRDye680LT dye with VISION using both the a-Si absorption filter and the ETFitc/Cy5 interference filter used in this work (Fig. S4c). Since the FOP trades off improved resolution for reduced collection efficiency, for a fair comparison of filter performance that does not include this tradeoff, a 250μm FOP is used with both filters. The dilution series is prepared by diluting a 12.5μM stock solution of IRDye680LT NHS ester (LI-COR) dissolved in 1x PBS by half until the concentration reaches 48.8nM. 35μL of each solution in the series is pipetted into chambered cover glass wells (103340, Grace Bio-Labs). Each solution is then imaged to produce an intensity vs concentration curve for both filters. By fitting a line to each curve and extracting the slope, we find that with the a-Si filter and the proposed optical frontend the sensor achieves sensitivities of 0.25mV/nM and 1.14mV/nM, respectively. This result indicates a 4.6x improvement in fluorescence collection efficiency with the proposed optical frontend, which corresponds closely with the theoretical results presented above.

The a-Si filter also passes more background light to the sensor. Firstly, the a-Si inherently has a long-pass characteristic as opposed the bandpass characteristic of the interference filter, such that additional out-of-band autofluorescence and ambient light is incident on the sensor. Secondly, compared to the proposed optical frontend, the a-Si filter provides significantly less excitation rejection. Images taken with both sensor of the excitation bleed-through from the 633nm laser operated 28mW/cm$^2$ (Fig. S4d), shows a more than 20x increase in measured excitation background with the a-Si filter. Increased background reduces image contrast and degrades SNR by increasing the shot noise in the sensor.

Moreover, the a-Si filter design is less versatile. In contrast to interference filters which are available with passbands across the entire visible and NIR spectra, the a-Si filter is only applicable with fluorophores that have emissions that fall in the 700-800nm region. While

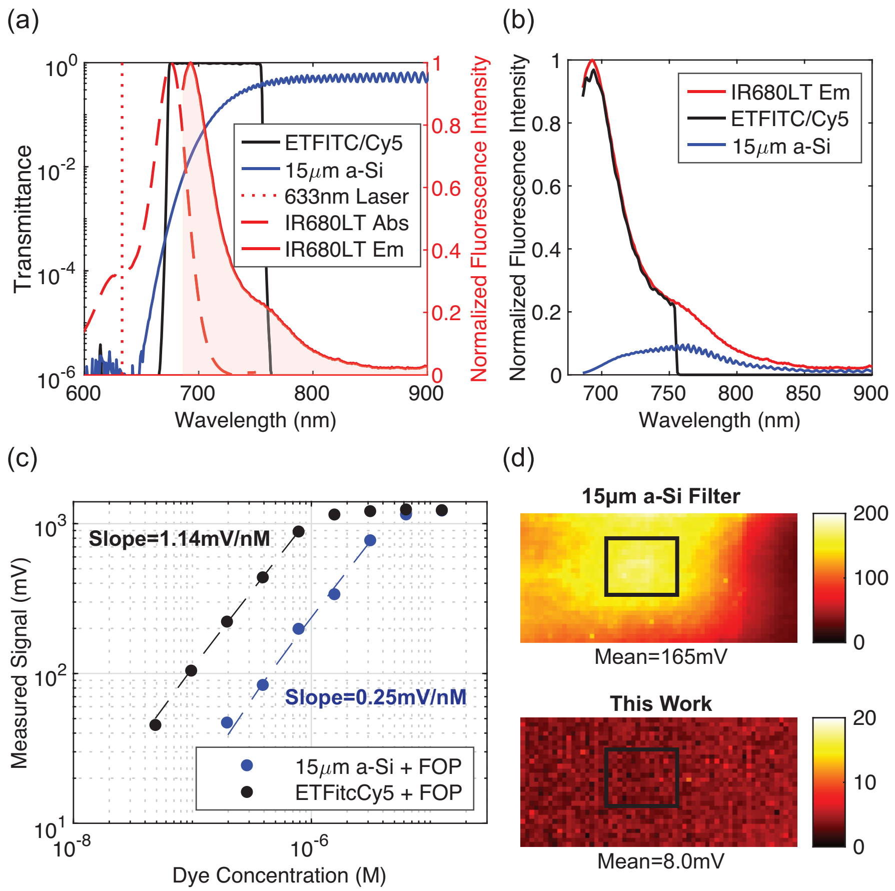


**Fig. S4. Comparison between the VISON optical frontend and the 15µm a-Si absorption filter.** (**a**) Transmittance spectra of both filters. (**b**) Plot of effective emission of IR680LT with each filter, which is the product of the filter transmittance and the emission spectra. (**c**) Plot of measured intensity across a 2x serial dilution of IRDye680LT NHS ester with sensors with both filters. By plotting the measured pixel intensity vs dye concentration for both sensors and extracting the slope from a linear fit, we see a 4.6x improvement in fluorescence collection efficiency with the VISION optical frontend. (**d**) Images of bleed-through from the 633nm laser used for excitation of IRDye680LT, operated at 28mW/cm$^2$ for both the 15µm a-Si filter and the proposed optical frontend.

different semiconductor materials can be used to cover other spectral regions, these often require different fabrication techniques that limit design flexibility. Furthermore, different absorption filters cannot be simply stacked for multiplexed imaging and, therefore, require complex pixel-level patterning of the filters across the sensor. As a result, multiplexed absorption-based filters represent an even further design and fabrication challenge. On the other hand, multi-bandpass interference filters that cover a variety of different fluorophore combinations with up to 4 distinct pass bands are widely available from commercial vendors.

#### *1.5 The effect of averaging on image quality*

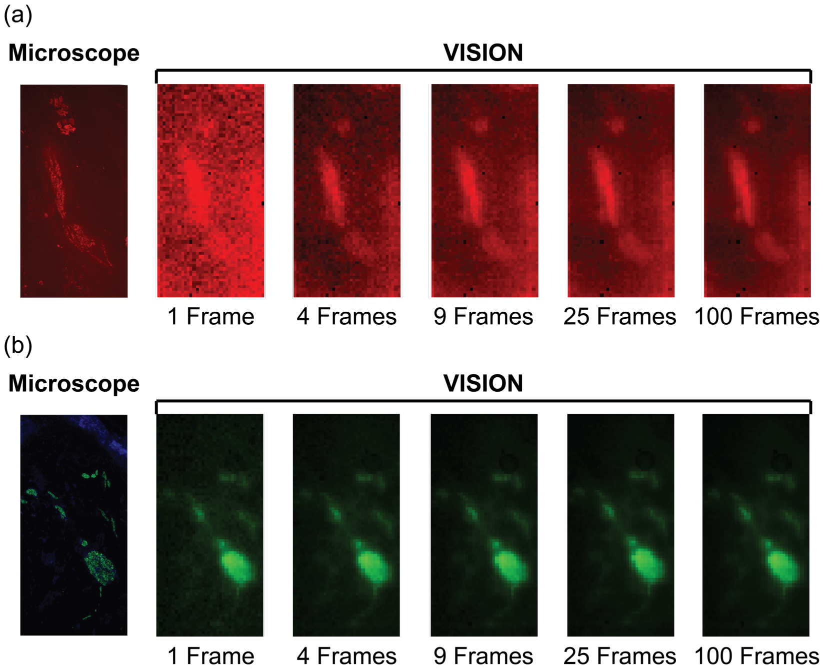


**Fig. S5. Effect of averaging on image quality.** (**a**) Images of the *ex vivo* prostate cancer sample in Fig. 6a.ii comparing a single frame from VISION using a 75ms exposure time to the same image after averaging 4, 9, 25, and 100 frames. The SNR of the image is linearly proportional to the square root of the number of averages. While a single frame is sufficient for detection, averaging significantly improves image quality. (**b**) A similar series of images showing the effects of averaging for the *ex vivo* nerve sample in Fig. 6a.v. The fluorescence in this channel is significantly brighter due to more optimal excitation near the absorption peak of the fluorophore, the higher quantum yield of the fluorophore, and properties of antibody used for staining. Since the SNR of a single frame is already high, the SNR improvement due to averaging is less noticeable.

### References

1. A. Schweikard, G. Glosser, M. Bodduluri, M. J. Murphy, and J. R. Adler, "Robotic Motion Compensation for Respiratory Movement during Radiosurgery," Computer Aided Surgery **5**, 263–277 (2000).
2. A. V Dsouza, H. Lin, E. R. Henderson, K. S. Samkoe, and B. W. Pogue, "Review of fluorescence guided surgery systems: identification of key performance capabilities beyond indocyanine green imaging," J. Biomed. Opt **21**, 80901 (2016).
3. E. P. Papageorgiou, B. E. Boser, and M. Anwar, "Chip-Scale Angle-Selective Imager for in Vivo Microscopic Cancer Detection," IEEE Trans Biomed Circuits Syst **14**, 91–103 (2020).
4. A. Gharia, E. P. Papageorgiou, S. Giverts, C. Park, and M. Anwar, "Signal to Noise Ratio as a Cross-Platform Metric for Intraoperative Fluorescence Imaging," Mol Imaging **19**, (2020).
5. E. P. Papageorgiou, H. Zhang, B. E. Boser, C. Park, and M. Anwar, "Angle-insensitive amorphous silicon optical filter for fluorescence contact imaging," Opt Lett **43**, 354 (2018).
